# ADH5-mediated NO Bioactivity Maintains Metabolic Homeostasis in Brown Adipose Tissue

**DOI:** 10.1101/2020.12.27.424489

**Authors:** Sara C. Sebag, Zeyuan Zhang, Qingwen Qian, Mark Li, Mikako Harata, Wenxian Li, Zhiyong Zhu, Leonid Zingman, Limin Liu, Vitor A. Lira, Matthew J. Potthoff, Alexander Bartelt, Ling Yang

## Abstract

Brown adipose tissue (BAT) thermogenic activity is tightly regulated by cellular redox status but the molecular mechanisms underlying this regulation are incompletely understood. Protein *S*-nitrosylation, the nitric oxide-mediated cysteine thiol modification of proteins, plays important roles in cellular redox regulation. Here we show that both diet-induced obesity (DIO) and acute cold exposure elevates protein S-nitrosylation of BAT proteins, including UCP1, to regulate thermogenesis. This effect in BAT is regulated largely by S-nitrosoglutathione reductase (GSNOR, ADH5), a denitrosylase that balances the intracellular nitroso-redox status. Loss of ADH5 specifically in BAT impairs UCP1-dependent thermogenesis during acute cold challenge and worsens metabolic dysfunction during diet-induced obesity. Mechanistically, we demonstrate that *Adh5* expression in BAT is controlled by the transcription factor heat shock factor 1 (HSF1) and administration of an HSF1 activator to the BAT of mice with DIO increased *Adh5* expression and significantly improved UCP1-mediated mitochondrial respiration. Together, these data demonstrate that ADH5 controls BAT nitroso-redox homeostasis to regulate adipose thermogenesis which may be therapeutically targeted to improve metabolic health.

**Highlights:**

- Thermogenesis induces protein S-nitrosylation modification in the BAT;
- ADH5, a major cellular denitrosylase, is required for maintaining BAT metabolic homeostasis under both overnutrition and cold stress conditions;
- Diet-induced obesity suppresses HSF1-mediated activation of *Adh5* in the BAT;
- ADH5 overexpression in BAT improves whole-body glucose homeostasis in obesity.

## INTRODUCTION

Brown adipose tissue (BAT) plays critical roles in converting intercellular and systemic lipid and glucose fuels directly into heat in response to thermogenetic challenge (Bartelt et al., 2011; Cannon and Nedergaard, 2004; Kajimura and Saito, 2014; Nedergaard et al., 2007; Saito et al., 2009; Townsend and Tseng, 2014). The BAT biology and physiology is regulated by a key signaling molecule, nitric oxide (NO), at multiple levels (Saha and Kuroshima, 2000). It has been demonstrated that NO is required for BAT thermogenic program by increasing blood flow following noradrenergic stimulation (Kikuchi-Utsumi et al., 2002; Nagashima et al., 1994), promoting BAT hyperplasia through alteration of the proliferation and differentiation of brown adipocyte precursors (Nisoli et al., 1998) and influencing brown adipocyte hypertrophy (Jankovic et al., 2017; Saha and Kuroshima, 2000). NO also modulates BAT lipid and glucose metabolism (Sansbury and Hill, 2014), sympathetic tone (Giordano et al., 2002; Nisoli et al., 1997) and mitochondrial biogenesis in response to acute thermogenic stimuli (Nisoli et al., 2003; Nisoli et al., 2004). Although basal level of NO is critical for maintaining cellular and systemic homeostasis, excess NO production contributes to the development of numerous pathological metabolic processes, including obesity (Chiarelli et al., 2000; Engeli et al., 2004; Jankovic et al., 2017; Noronha et al., 2005; Sansbury and Hill, 2014). However, the pathological role of NO bioactivity in the BAT in the context of obesity is largely unknown.

NO regulates cellular function largely through cGMP-dependent signaling, reactive oxygen and nitrogen species (ROS and RNS, respectively) signal transduction pathways events, and protein modifications (Stone and Marletta, 1996). In mice, diet-induced obesity (DIO) increases aberrant NO-mediated protein S-nitrosylation (SNO), leading to nitrosative stress that disrupts cellular homeostasis (Kaneki et al., 2007; Ovadia et al., 2011; Qian et al., 2018). We have recently shown that protein SNOs are elevated in the livers of obese patients compared to lean patients as well as in HFD-fed mice, resulting in impaired endoplasmic reticulum (ER) homeostasis and hepatic autophagy (Qian et al., 2019; Qian et al., 2018; Yang et al., 2015). In contrast, attenuating nitrosative stress by suppressing inducible nitric oxide synthase (iNOS) ameliorates obesity-associated insulin resistance in the liver, skeletal muscle and adipose tissue (Becerril et al., 2018; Perreault and Marette, 2001; Yang et al., 2015). With regards to pathophysiological studies in the BAT, it has been demonstrated that *in vitro*, excessive generation of NO inhibits lipolysis in rat brown preadipocytes (Nisoli et al., 1998); while in *vivo*, ablation of iNOS improves the energy balance of *ob/ob* mice in part by stimulating thermogenesis in brown fat cells (Becerril et al., 2010). Although these studies suggest a link between pathological NO bioactivity and impaired BAT function, the contribution of aberrant cellular nitrosative signaling in obesity-associated BAT metabolic dysfunction remains uncharacterized.

Alcohol dehydrogenase 5 (ADH5; also named S-nitrosoglutathione reductase, GSNOR) is the major cellular denitrosylase that regulates NO availability in the cell by catalyzing and regulating the breakdown of SNOs in order to balance the intracellular thiol redox status (Barnett and Buxton, 2017; Benhar et al., 2009; Liu et al., 2001). In plants, ADH5 plays a critical role in modulating tolerance to heat and cold stressors (Hussain et al., 2019; Leterrier et al., 2011; Lv et al., 2017). In mammals, dysregulation of ADH5 contributes to the pathogenesis of a diverse array of chronic diseases such as fatty liver disease, cardiovascular diseases and aging (Barnett and Buxton, 2017; Qian et al., 2018; Que et al., 2009; Rizza et al., 2018; Rizza and Filomeni, 2018; Sips et al., 2013; Tang et al., 2013). Notably, a recent study demonstrated that ADH5 regulates adipocyte differentiation (Cao et al., 2015). However, the role of ADH5-mediated protein denitrosylation during physiological cold adaptation or the pathogenesis of obesity-associated BAT dysfunction remains unknown.

In the present study, we demonstrate that ADH5 is required for maintaining BAT metabolic homeostasis by playing a previously unrecognized, yet key role, in balancing cellular NO bioactivity in brown adipocytes. Furthermore, we identified that dysregulation of the key proteostasis regulator heat shock factor 1 (HSF1a) leads to disruption of BAT ADH5 activation in the context of obesity. As such, targeting nitrosative stress in BAT by modulating the balance between ADH5 and iNOS actions may provide an avenue for the development of new therapies to improve metabolic health.

## RESULTS

### Obesity elevates protein S-nitrosylation and decreases cellular denitrosylation in the BAT

We previously reported that obesity induces nitrosative stress in the liver (Qian et al., 2018). To determine the impact of NO bioactivity on BAT function in response to various metabolic stressors, we first investigated the effects of nutritional overload on protein-S-nitrosylation (SNO) in the BAT using immunohistochemistry and a TMT-based S-nitrosylation Western blot analysis (Qu et al., 2014). Compared to mice fed a regular diet (RD), there was a significant increase in general protein SNO in the BAT from mice fed a HFD (Fig. 1A-C). Similarly, we found higher levels of BAT protein SNO in mice acclimated from thermoneutrality (30°C) to room temperature (22°C) and to acute cold (4°C) (Fig. 1D). To determine whether the elevation in BAT protein SNO was associated with an increase in cellular NO levels, we measured NO released by BAT explants isolated from mice fed with either a RD or HFD. Compared to RD mice, basal BAT NO production was elevated in mice fed a HFD (Fig. 1E), indicating obesity increases NO generation in the BAT. This obesity-associated elevation of NO is correlated with a reduction of UCP1-mediated mitochondrial respiration in the BAT from mice with DIO (Fig. 1F) using high resolution respirometry analysis.

**Figure 1.**
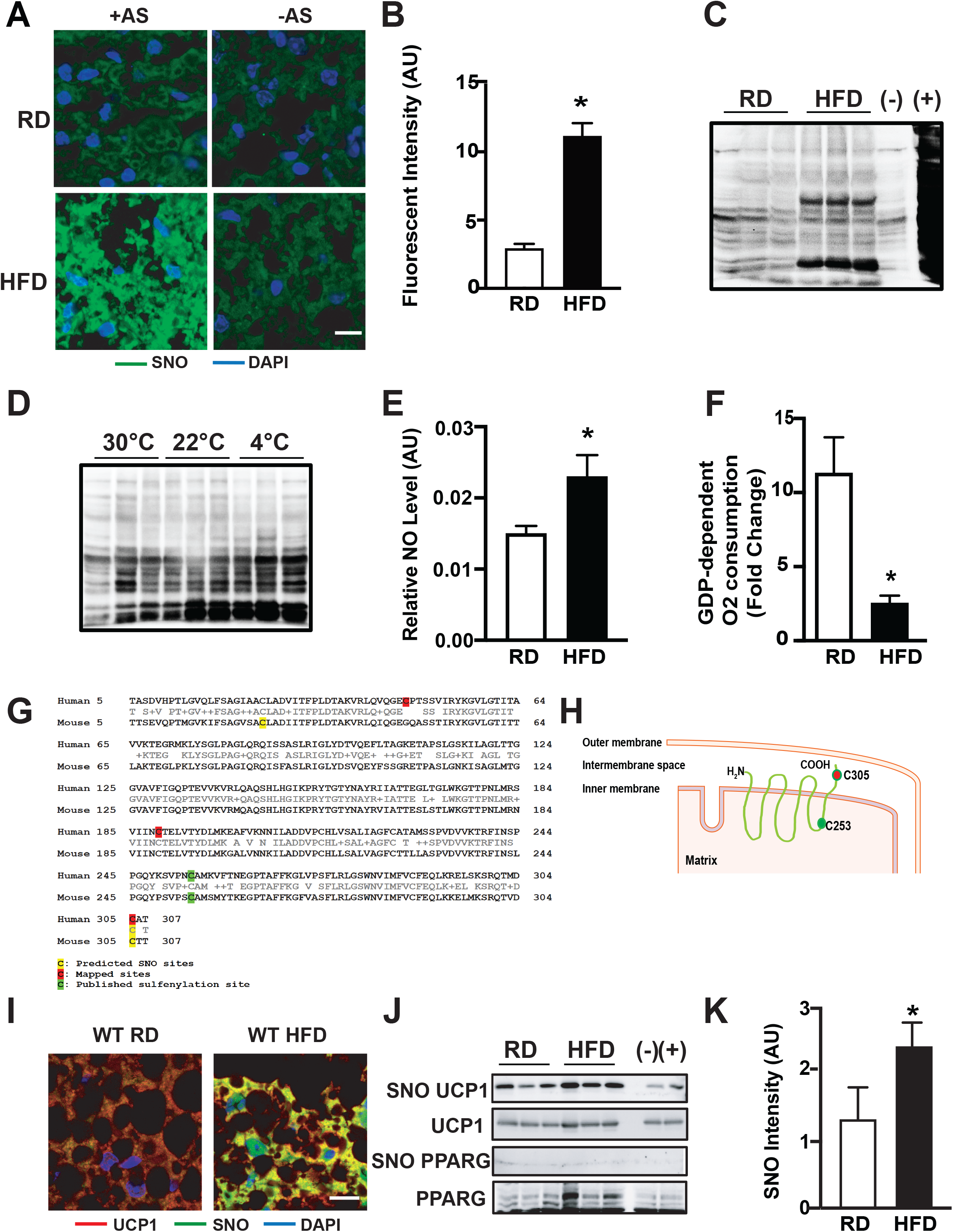
Diet-induced obesity elevates nitrosative stress in brown adipose tissue (BAT). **A&B**. Representative images (**A**) and quantification (**B**) of protein S-nitrosylation (SNO) staining in interscapular BAT (iBAT) from mice fed a regular diet (RD) or high-fat diet (HFD) for 10 weeks (63x). –AS: no ascorbate (negative control for S-nitrosylation staining). Scale bar: 10μm. Data are shown as Pearson’s correlation coefficient (n=3 mice/group). **C&D**. Western blots of Protein S-nitrosylation (SNO) in interscapular BAT (iBAT) from (C) mice fed a regular diet for 10 weeks; and (D) mice exposed to thermoneutrality (TN, 30°C for one week), room temperature (RT, 22°C) or cold (4°C for 24 hours), detected by iodoTMT switch assay (n=3 mice/group). (−): no ascorbate (negative control for S-nitrosylation detection); and (+): samples treated with SNAP, positive control for S-nitrosylation detection. **E**. Total nitric/nitrate NO release from supernatants of BAT explants isolated from mice on a RD (n=8 mice) or HFD (n=11 mice) for 10 weeks. **F**. High-resolution respirometry in fresh BAT from mice fed a RD or HFD for 10 weeks (n=4 mice/group). The UCP1-mediated respiration was calculated and reported as fold change of GDP-induced leak (1mM) over the baseline. **G&H**. Predicted SNO sites on mouse UCP1 and mapped SNO sites on human UCP1. **I**. Representative images (63X) of immunohistochemistry for S-nitrosylated UCP1 in the BAT from mice fed a RD or HFD for 10 weeks. Scale bar: 10μm. **J&K**. Representative western blots (H) and densitometric analysis (I) of S-nitrosylated proteins in the BAT from mice on a RD and HFD for 8 weeks. In western blots, each lane represents one individual mouse. All data are presented as means ± SEM. * indicates statistical significance compared to the RD groups, as determined by Student’s t-test (p<0.05).

Previous studies have shown that acute induction of thermogenesis in the BAT generates physiologic levels of mitochondrial ROS which in turn alters the redox status of the mitochondrial uncoupling protein 1 (UCP1), a key modulator of BAT thermogenic function (Chouchani et al., 2016). This redox-mediated modulation leads to activation of UCP1-dependent uncoupling (Chouchani et al., 2016). In contrast, studies have shown that excessive ROS and RNS impairs BAT mitochondrial function (Cui et al., 2019; Lee et al., 2020). GPS-SNO prediction tool (Xue et al., 2010) predicted that Cys25 and Cys305 are potential SNO sites on mouse UCP1, and cys305 as a potential SNO site on human UCP1 (Fig. 1G). We then mapped SNO sites on human UCP1 by mass spectra analysis using iodoTMT reagent, and found that Cys47, Cys189 and Cys305 are targeted by S-nitrosylation. Among these SNO sites, Cys305 is the common SNO site on both human and mouse UCP1(Fig. 1G&H). We next determined whether UCP1 is targeted by SNO in mouse BAT in vivo using immunohistochemistry and biotin switch analyses. As shown in Fig. 1I-K, UCP1 is not only targeted by S-nitrosylation modification, SNO-UCP1 expression was also significantly increased in the BAT of mice fed a HFD compared to RD controls. However, we did not detect SNO of PPARγ, an SNO target reported in mesenchymal stem cells (MSCs) (Cao et al., 2015), in BAT from either lean or obese mice (Fig. 1J). Collectively, these data indicate a potential role for protein SNO in the pathogenesis of obesity-mediated BAT dysfunction.

### ADH5 protects BAT against obesity-associated metabolic dysfunction

Alcohol dehydrogenase 5 (ADH5) is a major cellular denitrosylase that modulates NO availability in the cell by catalyzing the breakdown of SNOs (Barnett and Buxton, 2017; Benhar et al., 2009; Liu et al., 2004). To determine whether ADH5 is regulated by different metabolic stressors in the BAT, we first challenged WT RD mice with acute cold temperature (4°C) for 24 hours. As shown in Fig. 2A, cold exposure led to a significant increase in expression of ADH5 in the BAT. In contrast, in the context of obesity, ADH5 transcript and protein expressions were significantly reduced in the BAT (Fig. 2B&C). These data are consistent with a recent study demonstrating that obesity reduces cellular antioxidant capacity in the BAT (Alcala et al., 2017), while also demonstrating the stress-induced ADH5 regulation in BAT is context-dependent.

**Figure 2.**
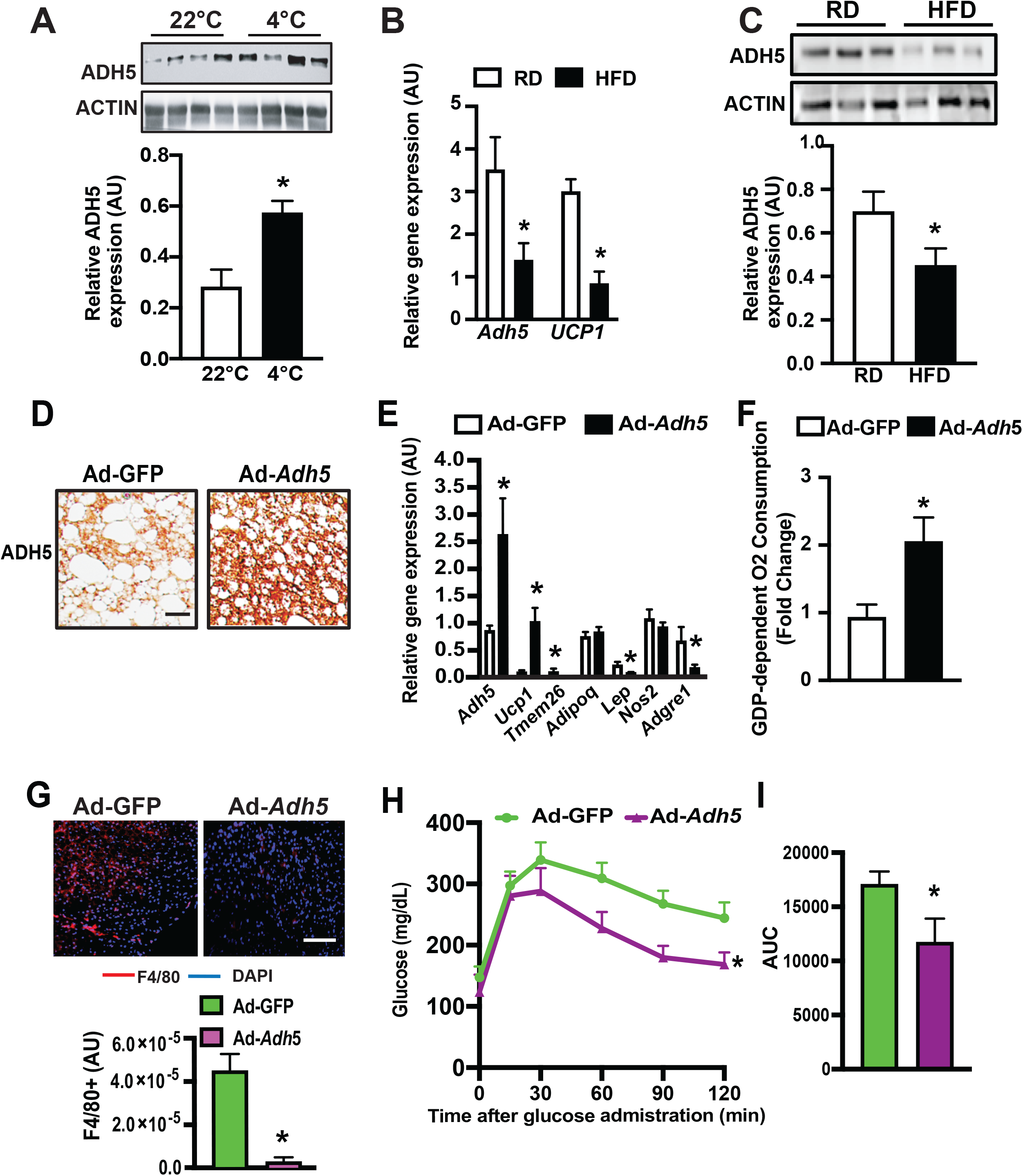
ADH5 protects metabolic dysfunction of BAT in the context of obesity. **A**. Representative western blots and densitometric analysis of tested proteins in the BAT from mice exposed to 22° C or 4° C for 24 hours. Data were normalized to expression of ACTIN (n=4 mice/group). **B**. Levels of mRNAs encoding the indicated genes in BAT from mice fed a RD (n=3 mice) or HFD for 10 weeks (n=5 mice), data were normalized to expression of *Hprt*. **C**. Representative western blots and densitometric analysis of ADH5 expression in BAT from mice fed a RD or HFD for 10 weeks (n=9-10 mice/group). Data were normalized to expression of ACTIN. **D**. Representative images (20X) of immunohistochemistry of ADH5 in the BAT from mice on a HFD for 8 weeks followed by interscapular transduction of Adeno-GFP (n=7 mice) or Adeno-*Adh5* (n=5 mice) (2.5×10^9^ pfu/mouse) for an additional 2 weeks. Scale bar: 10μm. **E**. Levels of mRNAs encoding the indicated genes; and **F**. GDP-dependent O_2_ consumption rates in BAT from mice in (D). Gene expression was normalized to *Gapdh*, n= 4 mice/group. **G**. Representative images (20X) and quantification of F4/80 staining in BAT from mice in (D). n=5-7 mice/group, 3-5 fields/replicate. Scale bar: 10μm. **H**. Glucose tolerance tests (GTT) and **I**. AUC in mice in (D). All data are presented as means ± SEM. * indicates statistical significance compared to 22° C in A, compared to the RD groups in B&C, and to the Adeno-GFP group in E-I, as determined by Student’s *t* test; p<0.05.

We next sought to determine whether gain of ADH5 function could improve BAT function in the context of obesity. For this purpose, we overexpressed *Adh5* in the BAT of adult mice using an adenovirus-mediated gene delivery approach via interscapular BAT injection (Balkow et al., 2016; Bartelt et al., 2018). First, we confirmed that ADH5 was overexpressed in the interscapular BAT (Fig. 2D&E). We then examined the effects of overexpression of *Adh5* on BAT function in the context of obesity. As shown in Fig. 2E, overexpression of ADH5 increased *Ucp1* while significantly reduced expression of the tested adipogenic markers in the BAT from mice on HFD (Fig. 2E). Furthermore, high resolution respirometry showed that these effects were concomitant with a increases in UCP1-mediated mitochondrial respiration in the BAT from mice on HFD (SFig. 1A and Fig. 2F).

Although the immunological properties of BAT are distinct from WAT and are comparatively less pronounced during the early stage of obesity (van den Berg et al., 2017; Villarroya et al., 2018a; Villarroya et al., 2018b), pathological inflammatory signals derived from both brown adipocytes and BAT-resident immune cells have deleterious effects on BAT recruitment and activation (Cao et al., 2018). We found that overexpression of *Adh5* significantly reduced the macrophagic marker *Adgre1* (F4/80) (Fig. 2E&G), which was associated with improved obesity-associated glucose intolerance (Fig. 2H&I). Together, these data show a protective role for ADH5 against obesity-associated BAT dysfunction.

### ADH5-deficency impairs metabolic function of BAT

To establish a BAT-specific ADH5 role, we crossed *Adh5*^fl^ mice (WT) (Wei et al., 2011) with a *Ucp1*-cre transgenic mouse line to generate BAT ADH5 KO (*Adh5*^BKO^) mice. *Adh5*^BKO^ mice had comparable core body temperature with *Adh5*^fl^ mice when housed at either room temperature (RT) or thermoneutrality (TN) (SFig. 1B). In addition, *Adh5*^BKO^ and *Adh5*^fl^ mice housed at RT had similar body composition (SFig.1B), and whole-body metabolic profiles as measured by comprehensive lab animal monitoring system (CLAMS; SFig. 1C-E). However, BAT from *Adh5*^BKO^ lean mice displayed a “white-like” phenotype at RT compared to littermate controls (Fig. 3A). Deletion of *Adh5* in the BAT increased expression of proinflammatory markers (*Nos2 and Adgre1*). In contrast, *Adh5* BAT-deletion reduced the brown and beige adipose identity markers (*Ucp1* and *Tmem26)*, but increased *Lep* transcript (Fig. 3B). Furthermore, *Adh5* deletion in the BAT significantly suppressed UCP1 protein expression (Fig. 3C&D) which was associated with elevated NO production and SNO of UCP1 (Fig. 3E&F).

**Figure 3.**
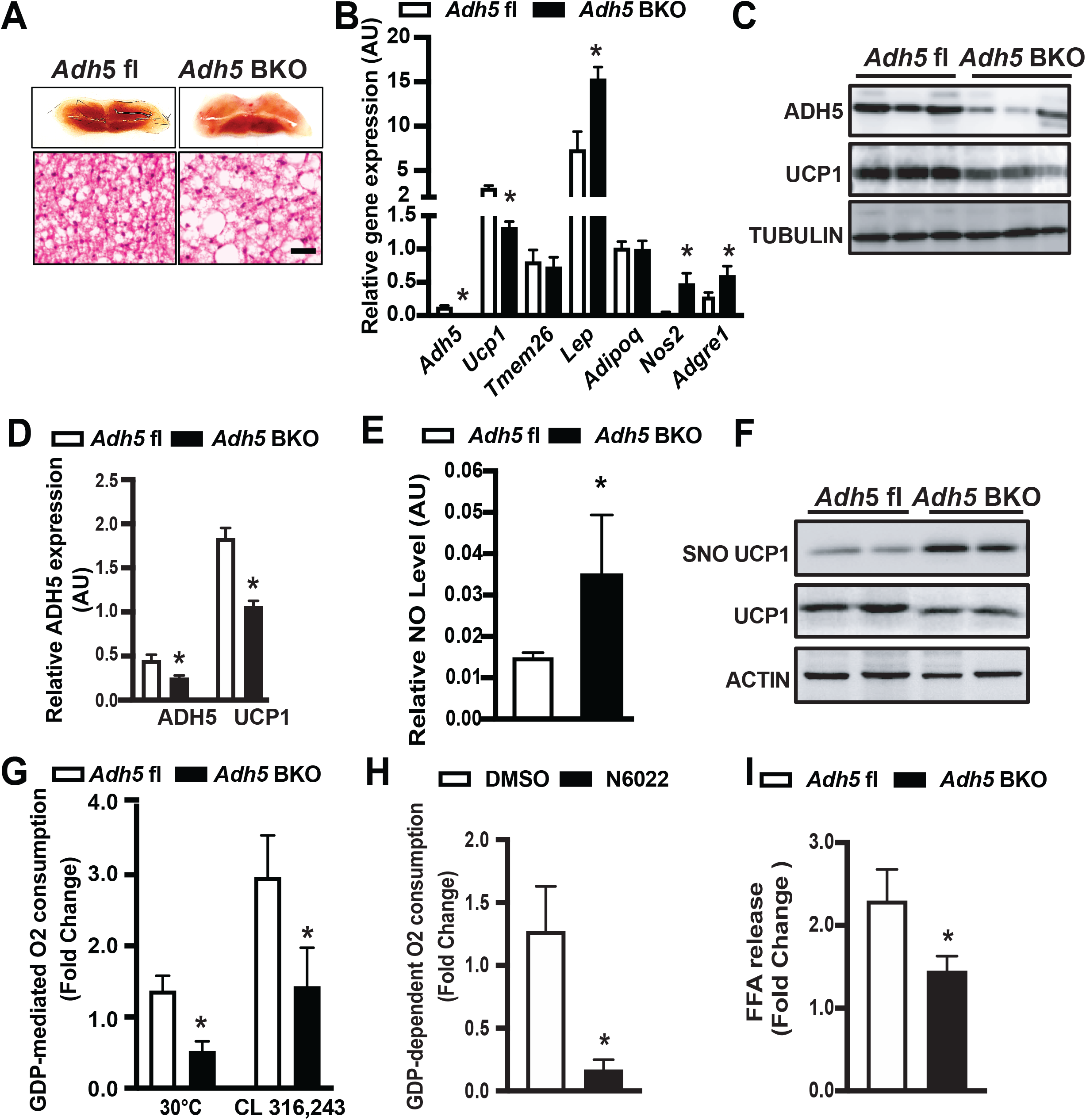
Deletion of *Adh5* impairs metabolic function of BAT. **A**. Representative light microscopy, and H&E images (10X) of BAT from Adh5^fl^ or Adh5^BKO^ mice raised at 22°C. **B**. Levels of mRNAs encoding the indicated genes in BAT from mice in (A). Expression is normalized to *Gapdh*; n=3-5 mice/group. **C**. Representative western blots of ADH5 and UCP1 expression in BAT from *Adh5*^fl^ and *Adh5*^BKO^ mice. **D**. Densitometric analysis is shown under the blots. Expression is normalized to that of TUBULIN; N=4-5 mice/group. **E**. Total nitric/nitrate NO release from supernatants of BAT explants from Adh5^fl^ or Adh5^BKO^ mice raised at 22°C. n= 3-8 mice/group. **F**. Representative western blots of S-nitrosylated UCP1 in the BAT from mice in (A). **G**. GDP-dependent O_2_ consumption in BAT from *Adh5*^fl^ and *Adh5*^BKO^ mice maintained at 30ºC (n= 3-4 mice/group), or injected with the β3-AR agonist CL316,243 (0.5mg/kg; n=4-5 mice/group); in **H**. BAT from wild type mice maintained at 30ºC following treatment with the ADH5 inhibitor, N6022 (20μM for 24 hours) or vehicle (DMSO, 0.2%); n=5 mice/group. **I**. Free fatty acid release measured in BAT explants from *Adh5*^fl^ and *Adh5*^BKO^ mice exposed to DMSO (0.1%) or isoproterenol (ISO, 1μM; 1hour). Data were normalized to protein concentration; n=5-6 mice/group. All data are presented as means ± SEM; *indicates statistical significance compared to the *Adh5*^fl^ groups in B, D, E, G, and I; and to the vehicle group in H, as determined by Student’s *t* test, p<0.05.

At thermoneutral temperatures, BAT activity is greatly reduced compared to standard RT (Cannon and Nedergaard, 2011). Therefore, to assess the contribution of ADH5 on UCP1-dependent mitochondrial respiration, we first performed mitochondrial respiration on fresh BAT excised from mice raised at TN. As shown in Fig. 3G, GDP-dependent, UCP1-mediated mitochondrial respiration was lower in BAT from *Adh5*^BKO^ mice compared to *Adh5*^fl^ littermate controls. BAT thermogenesis is activated via the β3-adrenergic pathway (Collins, 2011). Therefore, to determine if the effect of ADH5 deletion on UCP1-mediated thermogenesis is in part through reduction in β-adrenergic signaling, we further measured GDP-sensitive mitochondrial respiration in BAT fat pads from *Adh5*^BKO^ mice and Adh5^fl^ littermate controls that were treated intraperitoneally (i.p.) with CL316,243, an adipocyte-specific β3-adrenergic agonist (Cannon and Nedergaard, 2011). As shown in Fig. 3G, *Adh5-*deletion impaired β3AR-induced mitochondrial respiration as well. To confirm whether reduced mitochondrial respiration was attributed to defective ADH5 enzymatic activity, we excised BAT fat pads from WT mice fed with regular chow (housed at TN) and treated them *ex vivo* with an ADH5 chemical inhibitor, N6022 (Blonder et al., 2014). As shown in Fig. 3H, pharmacological inhibition of ADH5 significantly decreased GDP-dependent, UCP1-mediated mitochondrial respiration. Finally, lipolysis induced by the nonselective β-agonist isoproterenol (ISO) was also significantly reduced in BAT explants from *Adh5*^BKO^ mice compared to WT mice (Fig. 3I).

Previously, Cao et al. demonstrated that Adh5-deficient mesenchymal stem cells (MSCs) exhibited impaired adipogenesis (Cao et al., 2015). We then asked if ADH5 regulates BAT recruitment during thermogenesis by assessing brown preadipocyte differentiation using stromal vascular fraction (SVF) isolated from *Adh5*^fl^ mice followed by transduction of Adeno-Cre to delete *Adh*5. We found that, there were comparable expression levels of mature (*Pparγ* and *Ucp1)* and preadipocyte (*Pref-1*) markers, as well as lipid content between groups (SFig. 1F). Similar results were also observed in an immortalized human BAT cell line (hTERT-hA41BAT-SVF) (Xue et al., 2015) treated with the pharmacologic ADH5 inhibitor, N6022 (SFig. 1G&H). However, inhibition of ADH5 reduced oxygen consumption rate (OCR) in these cells (SFig. 1I), which is consistent with observation from Fig. 3H. Together, our data indicate a regulatory role for ADH5 in modulation of BAT activation.

### ADH5 is required for cold-induced thermogenesis in BAT

We found that ADH5 expression was increased by acute cold exposure in the interscapular BAT (Fig. 2A), the main organ that generates heat through non-shivering thermogenesis (Cannon and Nedergaard, 2004; Harms and Seale, 2013). Thus, to investigate the functional role of BAT-specific ADH5 in cold-induced thermogenesis, *Adh5*^fl^ and *Adh5*^BKO^ mice were weaned and raised at TN conditions and then challenged with acute cold exposure (4°C for 24 hours). Compared to the littermate wild type controls, both BAT and inguinal white adipose tissue (iWAT) from *Adh5*^BKO^ mice displayed an increased proportion of large-sized adipocytes (Fig. 4A). In addition, *Adh5*^BKO^ mice had significantly lowered core body temperature after either TN acclimation followed by cold exposure (Fig. 4B), or after activation of the β-adrenergic receptor (AR) by the β-adrenergic agonist norepinephrine (Fig. 4C). We then tested whether ADH5 in BAT is involved in the regulation of systemic thermogenic adaptation to cold exposure (4°C). Indirect mouse calorimetry analysis showed that cold exposure lowered both core and BAT temperature in *Adh5*^BKO^ mice compared to WT controls (Fig. 4D&E) which correlated with reduced whole-body oxygen consumption and energy expenditure (EE) (Fig. 4F&G). Furthermore, in line with our observations from mice housed at TN (Fig. 3F), *Adh5*^BKO^ mice showed a significantly reduced cold-induced GDP-dependent, UCP1-mediated mitochondrial respiration (Fig. 4H and SFig. 2A). Finally, we found compared to WT controls, that loss of BAT *Adh5* resulted in decreased expression of cold-induced genes that are normally activted as part of the thermogenic program (such as *Ucp1, Pparγ* and the cofactor *Ppargc1a*) (Fig.4I). Collectively, these results indicate that ADH5 is indispensable for adaptation of BAT to cold-induced thermogenesis.

**Figure 4.**
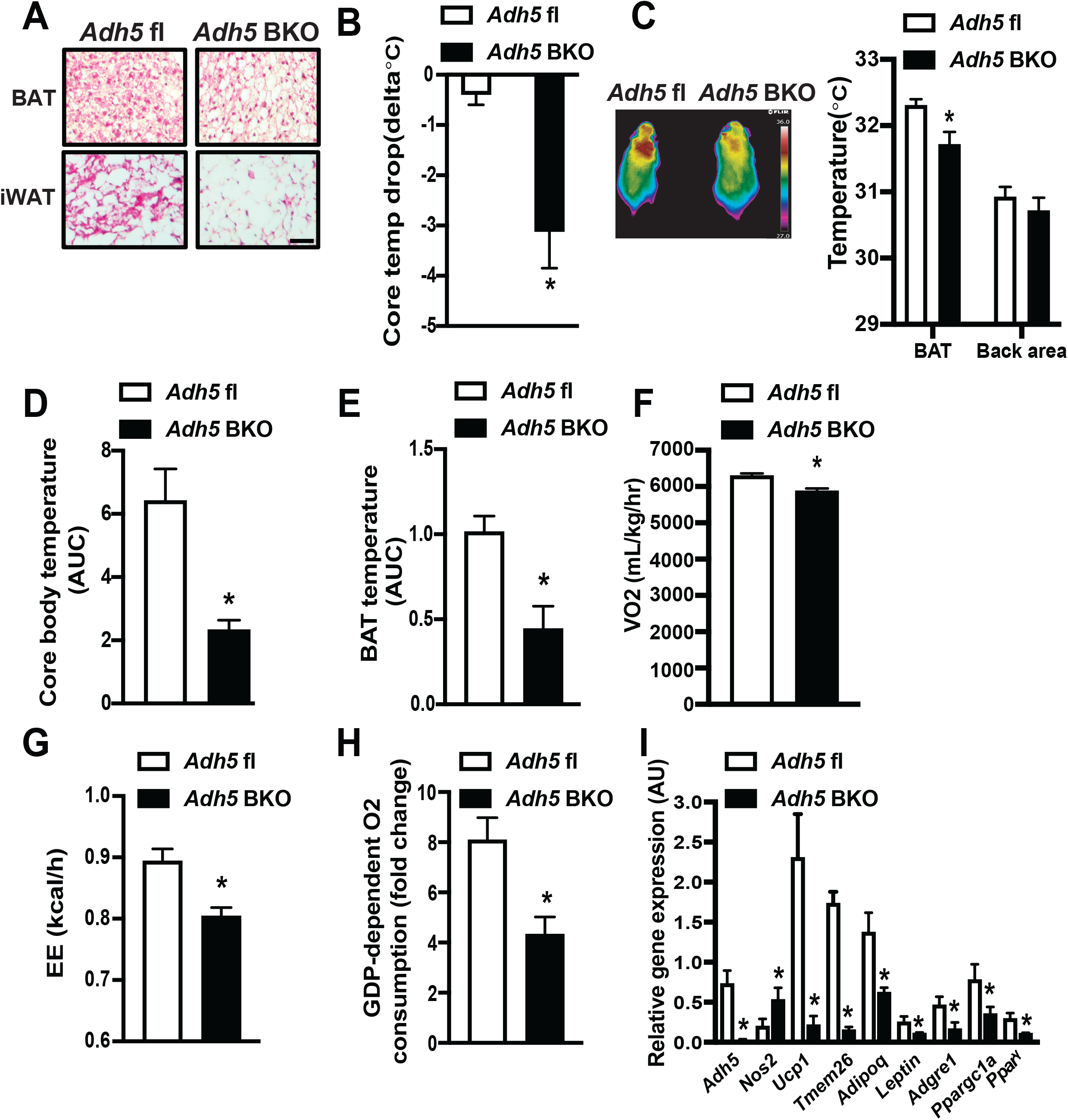
ADH5 is required for cold-induced BAT thermogenesis. **A**. Representative H&E images (10X) from BAT or iWAT; and **B**. Cold tolerance from *Adh5*^fl^ and *Adh5*^BKO^ mice housed at 30ºC following cold exposure to 4ºC for 24hr. Data were presented as difference in core body temperature between 30ºC and 4ºC. n=3-5 mice group. Scale bar: 100 μm. **C**. Representative high-resolution infrared images of body surface temperature of *Adh5*^fl^ and *Adh5*^BKO^ mice. Mice were injected with norepinephrine (0.3mg/kg) prior to imaging. Quantification of the images is shown by the side of image, n= 5 mice/group. **D**. Core temperature (AUC, area under the curve of dark cycles), **E**. BAT temperature (AUC of dark cycles), **F**. whole-body VO2, and **G**. whole-body energy expenditure (EE) measured in *Adh5*^fl^ and *Adh5*^BKO^ mice housed within a CLAMS at 4ºC (n=4-5 mice/group). **H**. GDP-dependent O_2_ consumption in BAT from *Adh5*^fl^ and *Adh5*^BKO^ mice (n=4/group) in (B). **I**. qPCR analysis measuring levels of mRNAs encoding the indicated genes in BAT from mice in (B). All data are presented as means ± SEM. *indicates statistical significance compared to the *Adh5*^fl^ group as determined by Student’s *t* test, p<0.05.

### Obesity impairs HSF1-mediated activation of *Adh5*

Obesity has been shown to suppress BAT metabolic function (Blondin et al., 2015; Himms-Hagen, 1985; Leitner et al., 2017; Shimizu et al., 2014; Shimizu and Walsh, 2015) while concomitantly increasing immune cell infiltration and ROS production (Alcala et al., 2017; Alcala et al., 2019). To establish the direct regulation of ADH5 in the context of obesity, we evaluated BAT function and systemic metabolic homeostasis in *Adh5*^BKO^and *Adh5*^fl^ mice fed a HFD. HFD-fed *Adh5*^BKO^ mice and *Adh5*^fl^ mice had comparable body weight and composition on the same diet (Fig. 5A). In addition, indirect mouse calorimetry demonstrated that activity, heat production and cumulative food intake were slightly decreased (yet did not reach statistical significance) in HFD-fed *Adh5*^BKO^ mice (SFig. 2B-D). However, *Adh5* deficiency in the BAT worsened obesity-associated glucose intolerance (Fig. 5B&C) and lowered whole-body oxygen consumption (Fig. 5D). At the tissue level, freshly isolated BAT from *Adh5*^BKO^ had a significant decrease in GDP-sensitive, UCP1-dependent mitochondrial respiration compared to explants from WT mice on a HFD (Fig. 5E).

**Figure 5.**
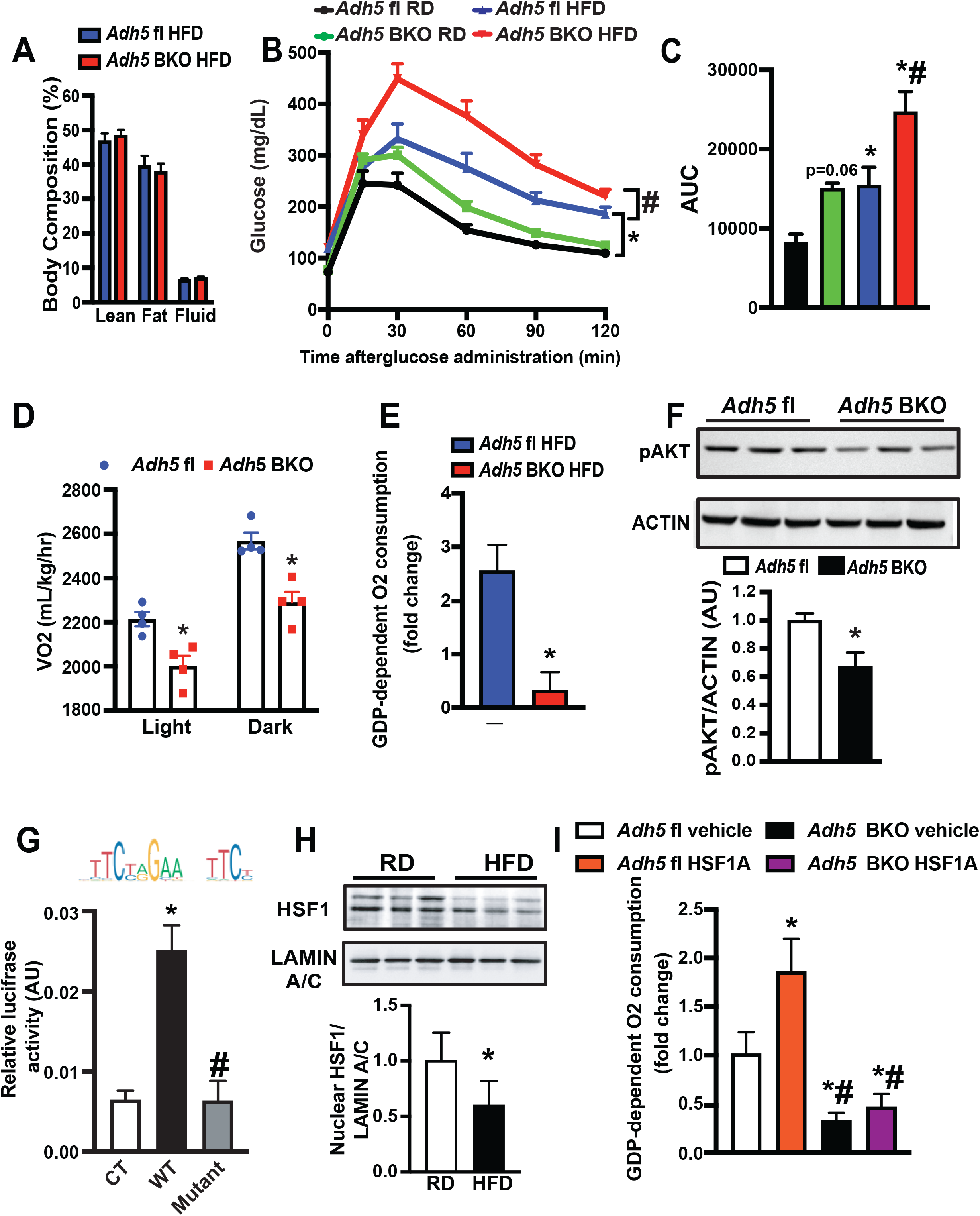
Obesity impairs HSF1-mediated *Adh*5 activation leading to BAT metabolic dysfunction. **A**. Body composition of *Adh5*^fl^ and *Adh5*^BKO^ mice on a HFD for 8 weeks, measured by nuclear magnetic resonance (NMR; n=7 mice/group). **B**. GTT and **C**. AUC from *Adh5*^fl^ and *Adh5*^BKO^ mice on a RD or HFD, n=11-12 mice/group. **D**. Whole-body VO2; and **E**. High-resolution respirometry analysis in BAT from *Adh5*^fl^ and *Adh5*^BKO^ mice fed a HFD. The UCP1-mediated respiration was calculated and reported as fold change of GDP-induced leak (1mM) over the baseline. n=4-5 mice/group. **F**. Insulin signaling in the BAT of *Adh5*^fl^ and *Adh5*^BKO^ mice on HFD as in (A). p-AKT: Akt^ser473^. Densitometric analysis is shown under the blots (n=3 mice/group). **G**. Activity of the *Adh5* promoter in HEK 293A cells 48 hours after transfection with the indicated constructs. Control cells (CT), PGL4.10 vector. Data were normalized to activity of Renilla Luciferase. Schematic representation of conserved HSF1 binding sequences identified in the promoter regions of *Adh5* using JASPAR is on the top of the panel. **H**. Representative western blots and densitometric analysis of HSF1 expression in nuclear fractions from BAT isolated from mice fed a RD or HFD for 16 weeks. n=6-8 mice/group. **I**. High-resolution respirometry analysis in BAT from WT mice on a HFD for 10 weeks followed by interscapular treatment with vehicle (DMSO; n=4 mice) or an HSF1A activator (1mM; n=7 mice) every other day for 2 weeks. For *Adh5*^BKO^, mice were fed on HFD for 10 weeks followed by DMSO or HSF1A activation (n = 7 mice/group). The UCP1-mediated respiration was calculated and reported as fold change of GDP-induced leak (1mM) over the baseline. All data are presented as means ± SEM. *indicates statistical significance compared to the *Adh5*^fl^ groups in C-E, to the CT in G, to the RD groups in H, and to the vehicle groups in I. # indicates genetic effects in mice with same treatment in H, and statistical significance compared to the WT in F. Statistical significance was determined by ANOVA in B, C, G, I; and Student’s *t* test in D-F, and H; p<0.05.

Malfunction of UCP1 and enhanced NO bioactivity activates innate immune cells in the BAT, leading to impaired glucose homeostasis (Bond et al., 2018). Therefore, we asked whether the loss of ADH5 function affected BAT immune balance. Indeed, we found a significant increase in HFD-mediated recruitment of F4/80 positive immune cells (SFig. 2E&F) which was concomitant with a significant increase in transcripts of the adipogenic markers (*Lep and Adipoq*), as well as the proinflammatory mediators (*Tnf* and *Il-6)*, and the macrophagic marker *Adgre1* (SFig. 2G). These changes in inflammatory markers were concomitant with worsened insulin signaling in the BAT from *Adh5*^BKO^ mice (Fig. 5F). These data provide evidence that ADH5-brown adipocyte expression is pivotal in preserving a healthy immune-metabolic balance of BAT during DIO.

We found that DIO suppressed *Adh5* expression in the BAT, therefore we next sought to determine the upstream signals involved in this suppression. The heat shock response (HSP) is a highly conserved physiological stress response to diverse stressors including cold-induced adrenergic activation of BAT (Matz et al., 1996). In an *in silico* analysis we found that HSF1 possesses binding elements at the *Adh5* promoter (Fig. 5G). To determine if HSF1 activates *Adh5, we* generated *Adh5* luciferase reporter constructs with HSF1 occupancy sites (−1683 to −1671) as well as a control construct lacking this binding site. HSF1 activated the *Adh5* promoter and this activation was abolished in cells expressing a mutant construct lacking this potential site (Fig. 5G). We then examined the nuclear expression of HSF1 in BAT from RD and HFD mice, and showed that DIO suppressed HSF1 nuclear localization in the BAT (Fig. 5H).

To explore the therapeutic potential of increasing HSF1 activity in the BAT, we treated WT DIO mice with HSF1A, a HSF1 activator, via interscapular BAT injection. Pharmacologic activation of HSF1 in HFD mice increased expression of *Hsf1*, HSF1 target genes (*Hsp90, Dnajb1 and Hsp27)*, as well as *Adh5* and *Ucp1* transcripts in the BAT (SFig. 2H). This increased *Adh5* expression by HSF1 activation was associated with increased GDP-sensitive, UCP1-dependent mitochondrial respiration in BAT fat pads compared to controls (Fig. 5I). Furthermore, activation of mitochondrial respiration BAT by HSF1 in the context of obesity was not altered in *Adh5*^BKO^ mice (Fig. 5I), indicating that ADH5 is a downstream factor in the HSF signaling cascade. Together these data indicate that activation of the HSF1-ADH5 axis protects BAT against obesity-associated metabolic dysfunction.

### Inhibiting iNOS in BAT alleviates DIO-associated metabolic dysfunction

Excessive NO production by parenchymal cells is one of the key factors that drives immune cell infiltration and polarization in the context of obesity (Sansbury and Hill, 2014). We observed higher iNOS expression in BAT from mice on HFD, which is worsened by deletion of *Adh5* (Fig. 6A, SFig. 2G). To define the major cellular source of the iNOS induction, we isolated SVF from *Adh5*^BKO^ and *Adh5*^fl/fl^ mice fed a HFD. While transcript levels of *Adh5, Nos2* or *Tnf* in the SVF were comparable between genotypes, BAT-specific *Adh5* deletion significantly increased expression of these markers in the BA fraction (Fig. 6B&C), which was in agreement with a higher NO production released from the BA (Fig. 6D). To further examine the direct role of NO bioactivity on BAT metabolic homeostasis in the context of obesity, we injected 1400W, a specific iNOS inhibitor (Garvey et al., 1997) into DIO mice interscapularly. As shown in Fig. 6E, treatment with 1400W decreased HFD-mediated NO production from BAT fat pads from both *Adh5*^fl/fl^ and *Adh5*^BKO^ mice. At the cellular level, GDP-dependent, UCP1-mediated mitochondrial respiration in freshly isolated BAT fat pads of obese mice was improved by 1400W injection regardless of genotype (Fig. 6F). Gene expression analyses further supported this notion showing that 1400W treatment reduced proinflammatory markers, while *Ucp1* was significantly increased in BAT from both *Adh5*^fl/fl^ and *Adh5*^BKO^ mice, respectively, on HFD (Fig. 6G). Taken together, these data demonstrate ADH5 deficiency in BA propagates obesity-associated BAT metabolic dysfunction is in part through induction of iNOS-mediated NO signaling (Fig. 6H).

**Figure 6.**
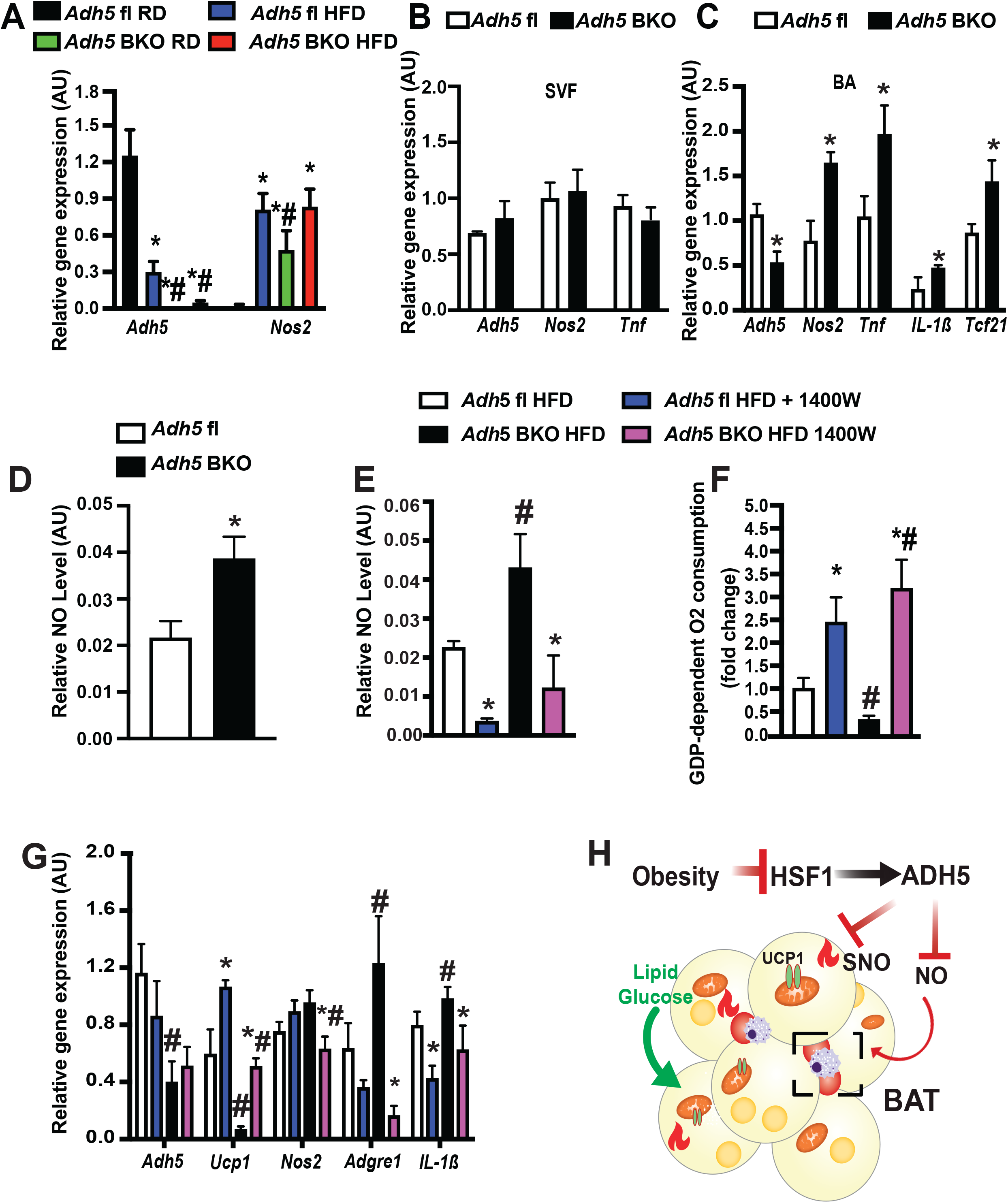
Inhibiting iNOS improves metabolic homeostasis in BAT from ADH5^BKO^ mice. **A**. qRT-PCR analysis measuring levels of mRNAs encoding the indicated genes in BAT from *Adh5*^fl^ and *Adh5*^BKO^ mice fed a RD or HFD for 16 weeks. n=4 mice/group. **B&C**. qRT-PCR analysis measuring levels of mRNAs encoding the indicated genes from mature brown adipocytes (B), and (C) SVF isolated from brown adipose tissues of *Adh5*^fl^ and *Adh5*^BKO^ mice (n=6 mice/group; 10wks HFD). **D**. Total nitric/nitrate NO release from mature brown adipocytes (C), and **E**. supernatants of BAT explants of HFD mice interscapularly injected with DMSO control or 1400W (0.2mg/kg) every 2 days for 3 weeks. n=7-12 mice/group. **F**. High-resolution respirometry analysis, and **G**. qRT-PCR analysis measuring levels mRNAs encoding the indicated genes in BAT from mice in (E). **H**. Model depicting ADH-mediated regulation of BAT metabolic function in the context of obesity. All data are presented as mean ± SEM. * indicates statistical significance compared to the *Adh5*^fl^-RD groups in A, to the *Adh5*^fl^ group in B-D, and indicates the treatment effects in mice with same genotype in E-G; # indicates genetic effects in mice with same treatments in A, and E-G. Statistical significance was determined by two-way ANOVA followed by Tukey’s multiple comparison in A, E-G; and Student’s *t* test in B-D; p<0.05.

## DISCUSSION

Intracellular Reactive Oxygen Species (ROS) and Reactive Nitrogen Species (RNS) are redox mediators that regulate diverse aspects of cellular processes and functions (Di Meo et al., 2016). Acute induction of thermogenesis in the BAT generates physiological levels of mitochondrial ROS which in turn alters UCP1-thiol redox status and activates UCP1-dependent uncoupling (Chouchani et al., 2016; Mailloux et al., 2012).

In contrast, under chronic metabolic-thermogenic conditions such as obesity, excessive levels of ROS are produced which promotes mitochondrial damage in the BAT (Cui et al., 2019; Kazak et al., 2017; Lee et al., 2020),. However, it is largely unknown whether and how NO bioactivity, another key redox mediator, modulates BAT function in health and disease. Our present study found that the protein S-nitrosylation signaling cascade plays a key role in regulation of BAT thermogenic function.

NO bioactivity is tightly regulated by the balance of NO availability and the capacity to remove it in part by denitrosylases(Benhar et al., 2009). For the former, two NOS isoforms – eNOS and iNOS, are expressed in the BAT (Engeli et al., 2004; Kikuchi-Utsumi et al., 2002) with temporal studies revealing that eNOS is expressed early during cold acclimation whereas iNOS is elevated in the late phase (Otasevic et al., 2011). On the other hand, ADH5 (Cao et al., 2015) and thioredoxin reductase 1 (Ayyappan and Nagajyothi, 2017), both of which serve to denitrosylate and modify protein function (Nakamura and Lipton, 2016), are expressed in BAT. These data suggest that within the BAT there exists a fine-tuned equilibrium between protein *S*-nitrosylation by NOS synthases and denitrosylases. However, the pathophysiological role of S-nitrosylative signaling in brown adipose tissue has yet to be well defined. Our study provides the first evidence that DIO increases BAT NO production and elevates general protein S-nitrosylation, including UCP1, in the BAT. On the other hand, we found that the key cellular denitrosylase, ADH5 protects against impairment of UCP1-mitochondrial respiration and inflammation as well as glucose intolerance in obese mice.

Previous studies investigating the modulation of nitrosative signaling in the adipose tissue have involved deletion or inhibition of NOS expression and activity (Becerril et al., 2018; Fujimoto et al., 2005; Tsuchiya et al., 2007; Zanotto et al., 2017), thereby diminishing all forms of NO bioactivity while also impairing both cGMP and S-nitrosylation-mediated signaling. Ultimately, these approaches to dissect the function of protein S-nitrosylation in regulating BAT physiology are very challenging. In order to investigate the distinct roles of protein S-nitrosylation in regulating BAT function, we focused our study on the key denitrosylase, ADH5. ADH5 has been shown to play an important role in multiple cellular responses, including β-AR signaling, endocytosis, inflammation, angiogenesis and apoptotic cell death (Barnett and Buxton, 2017). In particular, we investigated the hypothesis that ADH5 regulates BAT metabolic and thermogenic function by balancing intracellular NO bioactivity. We showed that BAT-specific loss of ADH5 aggravated obesity-mediated metabolic dysfunction and inflammation, as well as impaired UCP-1 mediated mitochondrial respiration. Furthermore, BAT-ADH5 deletion led to the dysregulation of thermogenic responses to both cold or direct β-AR3 activation.

In our study, we found that obesity promotes SNO of UCP1, which is augmented by *Adh5* deletion in the BAT. Using GPS SNO tool (Xue et al., 2010), we further predicted two potential SNO sites on the mouse UCP1 (Cys25 and Cys305), and mapped the SNO sites on human UCP1 (Fig. 1G). It is important to note that sulfenylation of UCP1 at Cys253 (localized on the matrix side of mitochondria) is required for BAT function during thermogenesis (Chouchani et al., 2017), however, future studies focusing on delineating fine-tuned UCP1 modifications by redox cues generated from acute and chronic stress may provide additional insights into the thermogenic function of BAT in health and disease. We do recognize that the role of ADH5 in regulating BAT metabolic function could be attributed to either UCP1-dependent uncoupling or a UCP1-independent action. In fact, we also observed that ADH5 expression positively correlates with *Ucp1* transcript. In addition, in other systems multiple thermogenic regulators have been demonstrated are tightly regulated by protein SNO. In the heart two key proteins involved in β2-AR receptor regulation, GRK2 and β-arr2, are shown to be targeted by SNO (Whalen et al., 2007). In skeletal muscle, the control of calcium-handling machinery is also governed by SNO, such that it has been shown that ADH5-KO mice exhibit an impaired inotropic response to isoproterenol in part by reducing denitrosylation of RyR2 which leads to pathological calcium leak (Beigi et al., 2012; Stomberski et al., 2019). Of note, a recent study demonstrated that SNO of the peroxisome proliferator-activated receptor gamma (PPARγ) in MSCs reduces adipogenesis while increasing osteoblastogenesis (Cao et al., 2015). In our study, we did not observe detectable SNO of PPARγ in the BAT, nor an alteration of Adh5 deletion on BA differentiation (Fig.1G & SFig.1E-J). We postulate that this regulation might be context-dependent and is an outcome of whole-body deletion vs BAT-specific deletion.

Recently, cellular proteostasis has been implicated as a key aspect of BAT cold adaptation (Bartelt et al., 2018). Here we showed that both acute cold acclimation and obesity induces protein S-nitrosylation in the BAT (Fig 1.D). It is possible that, during cold exposure and nutritional overload, damaged proteins are targeted by S-nitrosylation modification for further processing, and the required resolution is regulated in part through ADH5-mediated signaling. Mechanistically, our study demonstrates that ADH5 is directly induced by HSF1. HSF1, a master regulator of the HSR, is classically induced by heat shock, but is also activated by cold challenge (Ma et al., 2015; Reinke et al., 2008; Sarge and Cullen, 1997). Here, we showed that loss of the HSF1-ADH5 signaling cascade contributes to the metabolic dysfunction of BAT in mice with DIO. This is in line with previous studies that showed treatment with celastrol activates HSF1 leading to improved metabolic function in the BAT of obese mice (Ma et al., 2015). Herein, our study provides mechanistic insight into how the HSF1-mediated transcriptional program controls cellular nitro-redox signaling by modulating expression of *Adh5*. Ultimately, future studies defining the dynamic regulation of BAT-ADH5-HSF1 may provide a potential therapeutic target in the context of obesity-associated metabolic disease.

Another key feature of obesity-associated metabolic abnormalities is low grade chronic inflammation in the adipose tissue, contributing to insulin resistance and metabolic abnormality (Hotamisligil, 2006; Xu et al., 2003). Prior studies have demonstrated that pathological inflammatory signaling derived from both brown adipocytes and BAT-resident immune cells have deleterious effects on BAT recruitment and activation (Bartelt et al., 2018; Cao et al., 2018). Moreover, in rodents, supplementing with anti-inflammatory molecules (*e*.*g*. epigallocatechin gallate, and berberine) increases BAT thermogenic activity (Okla et al., 2017). Excessive NO production by parenchymal cells is one of the key factors driving immune cell infiltration and polarization in the context of obesity (Sansbury and Hill, 2014). Our study showed that *Adh5* deletion increased *Nos2* expression in the BA (Fig. 6A-C) with concomitant elevation of BA-derived NO production as well as proinflammatory markers (Fig. 6C&D). We further found that these processes were attenuated by chemical inhibition of iNOS (via 1400W) in the BAT (Fig. 6D). Thus, an important question for future studies is to determine the impact of ADH5-mediated BA NO bioactivity on the immune signature, vascularization, sympathetic tone as well as remodeling in the BAT in health and disease.

Taken together, our study demonstrates that cellular nitrosative signaling is critical for BAT health. Identifying the molecular mechanisms underlying the immuno-metabolic function of the BAT through ADH5-mediated nitro-redox signaling will help establish a framework for developing novel therapies to ameliorate obesity-mediated BAT dysfunction.

## ACKNOWLEDGMENTS

We thank Drs. Huojun Cao, Renata P. Pereira and Julien Sebag at the University of Iowa and Drs. Ana Paula Arruda and Grace Yankun Lee at the Harvard T.H. Chan School of Public Health for technical supports. We are grateful to for Dr. Dale E. Abel at the University of Iowa for scientific discussions and insights.

## AUTHOR CONTRIBUTIONS

L.Y. and S.C.S. designed the study and wrote the manuscript. S.C.S., Q.Q., Z.Z., M.L., M.H., W.L., performed the experiments. V.L., L.L., M.J.P., and A.B. provided critical reagents and scientific suggestions on the manuscript. L.Y. conceived and supervised the study. Z.Y.Z. is supported by an American Heart Association Predoctoral Award (19PRE34380258); L.Y. is supported by an American Diabetes Association Innovative Basic Science Award (1-18-IBS-149) and NIH R01 DK108835-01A1.

## SUPPLEMENTAL FIGURES

**Supplemental Figure 1.**
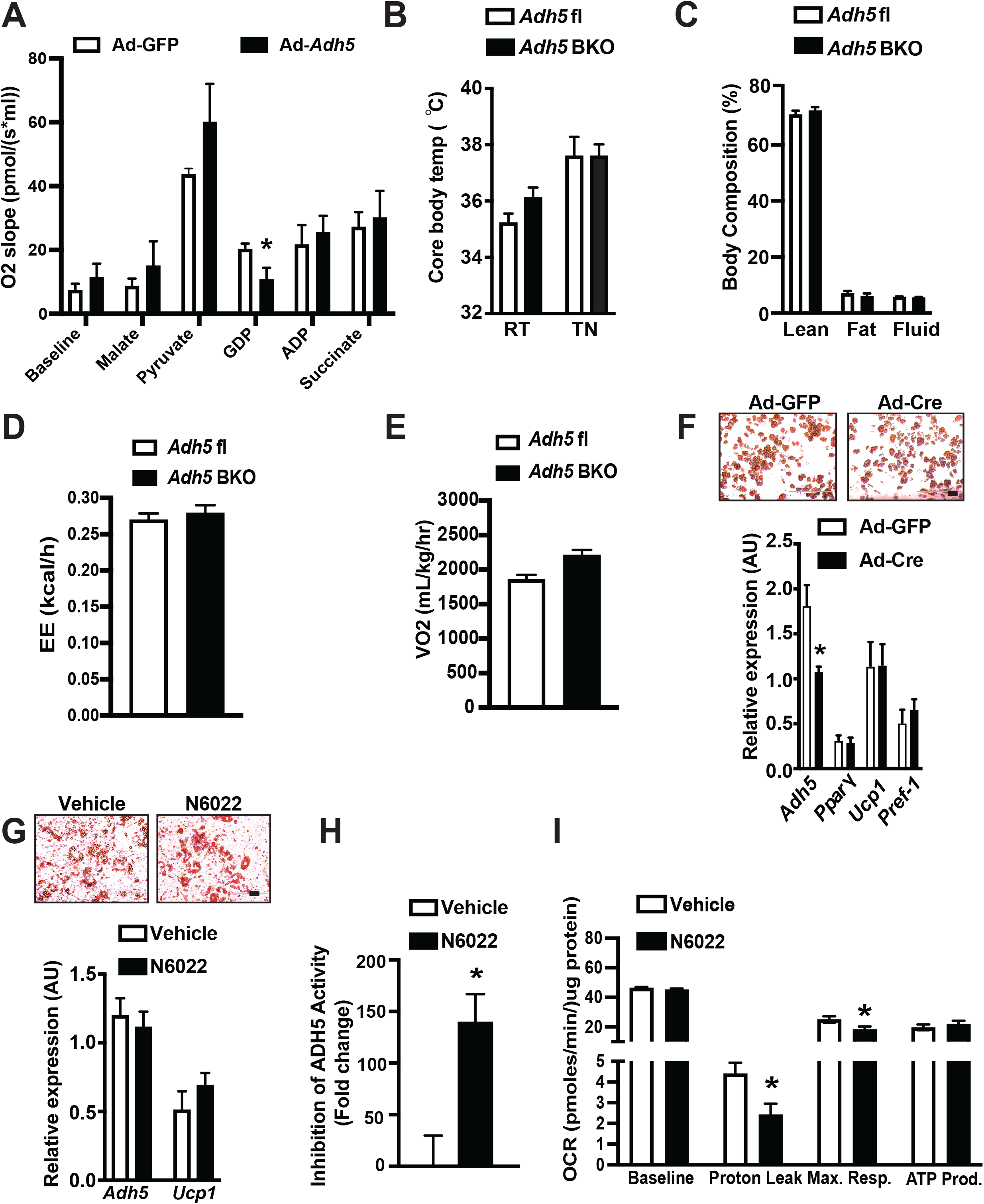
Metabolic profiles of mice with loss of ADH5 in the BAT. **A**. O_2_ consumption rates in fresh isolated BAT from mice on a HFD for 8 weeks followed by interscapular transduction of Adeno-GFP or Adeno-*Adh5* (n=4 mice/group). **B**. Core body temperature measured in *Adh5*^fl^ or *Adh5*^BKO^ mice housed at RT (22ºC) or TN (30ºC) for 3 weeks. n =5-11 mice/group. **C**. Body composition of *Adh5*^fl^ or *Adh5*^BKO^ mice on RD for 8 weeks. n=5-6 mice/group. **D**. Energy expenditure and **E**. whole body VO2 were measured in *Adh5*^fl^ or *Adh5*^BKO^ mice housed in CLAMs at RT, n= 3-4 mice/group. **F**. Representative Oil red O staining (10X), and qPCR analysis measuring levels of mRNAs encoding the indicated genes in BA differentiated from SVF of *Adh5*^fl^ mice. Preadipocytes were transduced with Adeno-GFP or Adeno-Cre (1×10^12^ pfu/mouse) at day 5 of differentiation. Data were normalized to *Gapdh* (n = 3, biological replicates). **G**. Representative Oil red O staining (10X), and qPCR analysis measuring levels of mRNAs encoding the indicated genes in differentiated, and **H**. ADH5 activity in human BAT cells (hTERT A41hBAT-SVF) in the presence or absence of 25 nM N6022. Gene expressions were normalized to 18S. n=3 experimental replicates. **I**. O_2_ consumption rates in cells in (**G**). All data are presented as means ± SEM; *indicates statistical significance as determined by Student’s *t* test, p<0.05.

**Supplemental Figure 2.**
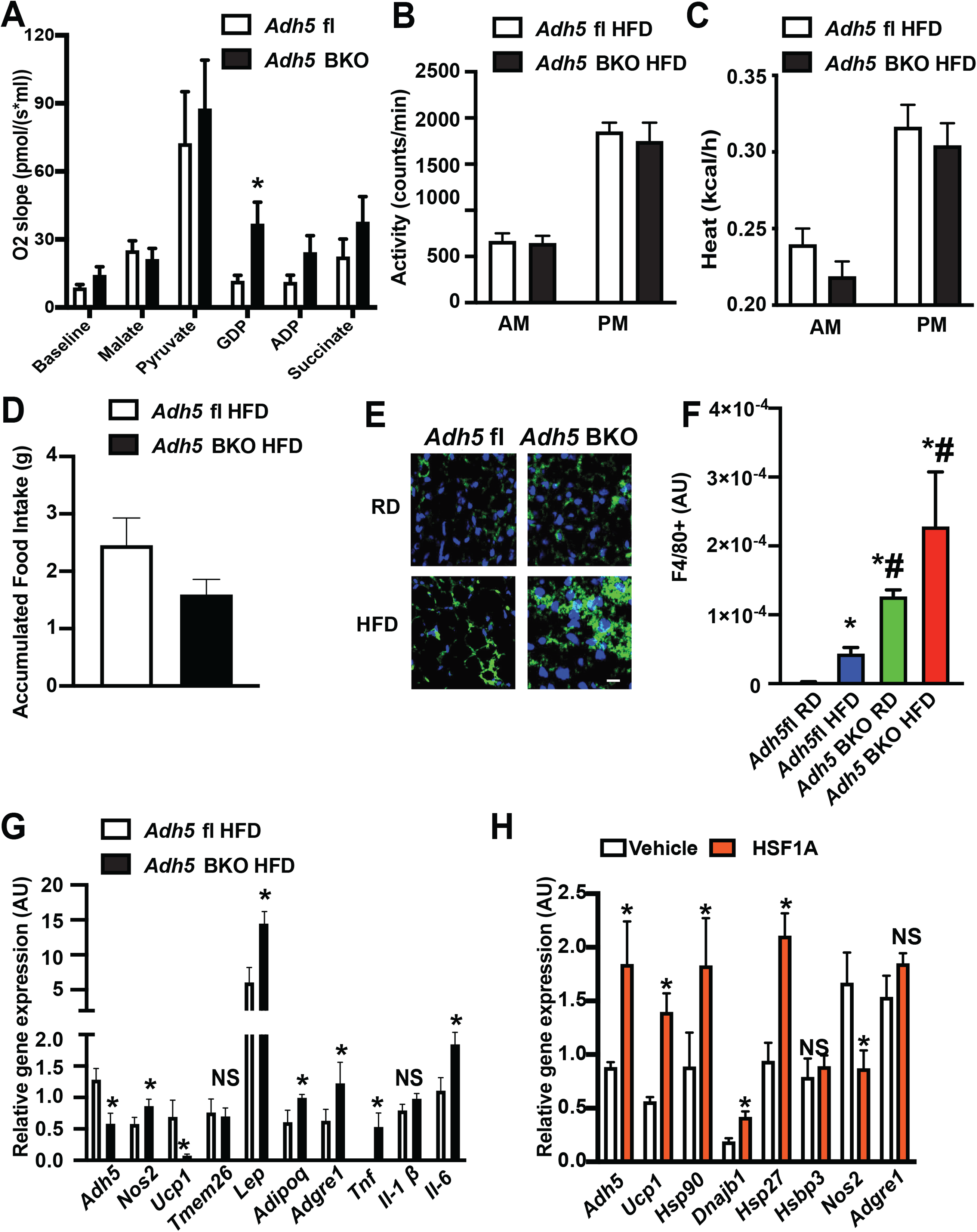
Inflammatory and metabolic profiles in BAT with loss of *Adh5* and chemical activation of ADH5. **A**. O_2_ consumption rates in fresh isolated BAT from *Adh5*^fl^ and *Adh5*^BKO^ mice housed at 30ºC following cold exposure to 4ºC for 24hr. n=4/group. **B-D**. Activity, heat and food intake were measured in HFD-fed (8 weeks) *Adh5*^fl^ or *Adh5*^BKO^ mice housed in CLAMs at RT, n=4-5 mice/group. **E&F**. Representative immunofluorescent (20X) staining and quantification of F4/80 in BAT from *Adh5*^fl^ (n=3-7 mice/group) and *Adh5*^BKO^ (n=3-5 mice/group) mice fed a RD or HFD for 16wk. Scale bar: 10μm. Quantification includes 3-5 fields/replicate. **G**. qPCR analysis measuring levels of mRNAs encoding the indicated genes in BAT from mice in (**B**) (n=3-5 mice/group). **H**. qPCR analysis measuring levels of mRNAs encoding the indicated genes in BAT from mice fed a HFD for 10 weeks followed by interscapular treatment with vehicle (DMSO; n=4 mice/group) or an HSF1A activator (1mM; n=7 mice/group) every other day for 2 weeks. All data are presented as means ± SEM; *indicates statistical significance as determined by ANOVA (**F**), or Student’s t-test (**A&G&H**). p<0.05.

## MATERIALS AND METHODS

### Mouse models

Animal care and experimental procedures were performed with approval from the University of Iowa’s Institutional Animal Care and Use Committee. C57BL/6J mice (The Jackson Laboratory, 000664), UCP1-cre (The Jackson Laboratory, B6.FVB-Tg(UCP1-cre)1Evdr/J, 024670), and *Adh5*^fl^ (Wei et al., 2011) mice were kept on a 12-hour light/dark cycle. Mice used to generate the DIO model were placed on a 60% kCal high-fat diet (Research Diets, D12492) immediately after weaning (i.e., at 3 weeks of age). Adenovirus-*Adh5(Qian et al*., *2018)*, and adenovirus-GFP were delivered to mice interscapularly at a titer of 2.5×10^9^ pfu/mouse(Bartelt et al., 2018). HSF1A (Millipore,1196723-93-9) was interscapularly administrated to mice at a dosage of 1mM/mouse every other day for 1 week. 1400W (Sigma, 214358-33-5) was interscapularly administrated to mice at 0.2mg/kg every other day for 2 weeks by sub-BAT injections. For cold-exposure and thermoneutrality experiments, mice were singly housed in an environmental chamber during a 12-hour light cycle (Metabowl; Jencons Scientific). Mice were given access to standard 2920X Teklad Global Diet and water *ad libitum* while in cages. All tissues were harvested, frozen in liquid nitrogen, and kept at −80°C until processed.

### Cell Culture

HEK 293A cells (ATCC) were cultured in DMEM medium with 10% cosmic calf serum (CCS) and 1% penicillin-streptomycin. Cells were transfected with DNA constructs (0.5 ug/well in a 24-well plate) using Lipofectamine 2000 reagent (Thermo Fisher Scientific, L3000015) in Opti-MEM medium (Thermo Fisher Scientific, 31985-062). At 48 hours post transfection, the activities of firefly luciferase and Renilla luciferase were measured using the Luciferase Assay System (Promega, E1500) and Renilla Luciferase Assay System (Promega, E2810), separately.

Primary brown adipocytes were isolated from iBAT depots of *Adh5*^fl^ mice as previously described (Gao et al., 2017; Ikeda et al., 2017). Briefly, fat tissues were minced and digested in 1mg/ml Liberase (Roche, 5401119001) in KRBH buffer in a 37^0^C water bath with agitation for 30 minutes, followed by filtration through nylon mesh. The samples were centrifuged at 500g for 10 minutes at 10ºC. The floating mature adipocytes were collected and cultured in DMEM/F-12 containing 10% cosmic calf serum (CCS) and 1% penicillin-streptomycin. The pellets were resuspended in DMEM/F-12 containing 10% calf serum (CCS) and 1% penicillin-streptomycin (basal growth media, BGM) and plated on 6 well plates. Cells were then grown for 5 days in BGM, infected in BGM containing Adeno-GFP (1×10^10^) or Adeno-ADH5 (1×10^12^) and differentiated in BGM containing 0.25nM 3-Isobutyl-1-methylxanthine (IBMX, Sigma, I5879), 1uM Rosiglitazone (Sigma, 557366-M), 0.125nM Indomethacin (Sigma, 405268), 1nM T3 (Sigma, T5516), Humulin R (Lily, Humulin R U-500, 0002-8501-01) and 2ug/ml dexamethasone (Sigma, D4902) for up to 10 days with media change every other day. The hTERT A41hBAT-SVF was obtained from ATCC (CRL-3385) and cells were grown in BGM medium, followed differentiation with or without 25nM N6022 (Sigma, 1208315-24-5).

### Quantitative Real-time RT-PCR

Total RNA was isolated using TRIzol reagent (Invitrogen,15-596-018) and reverse transcribed into cDNA using the iScript cDNA synthesis kit (Bio-Rad, 1708890). Quantitative real-time RT-PCR analysis was performed using SYBR Green (Invitrogen, KCQS00). Primer sequences are as follows:

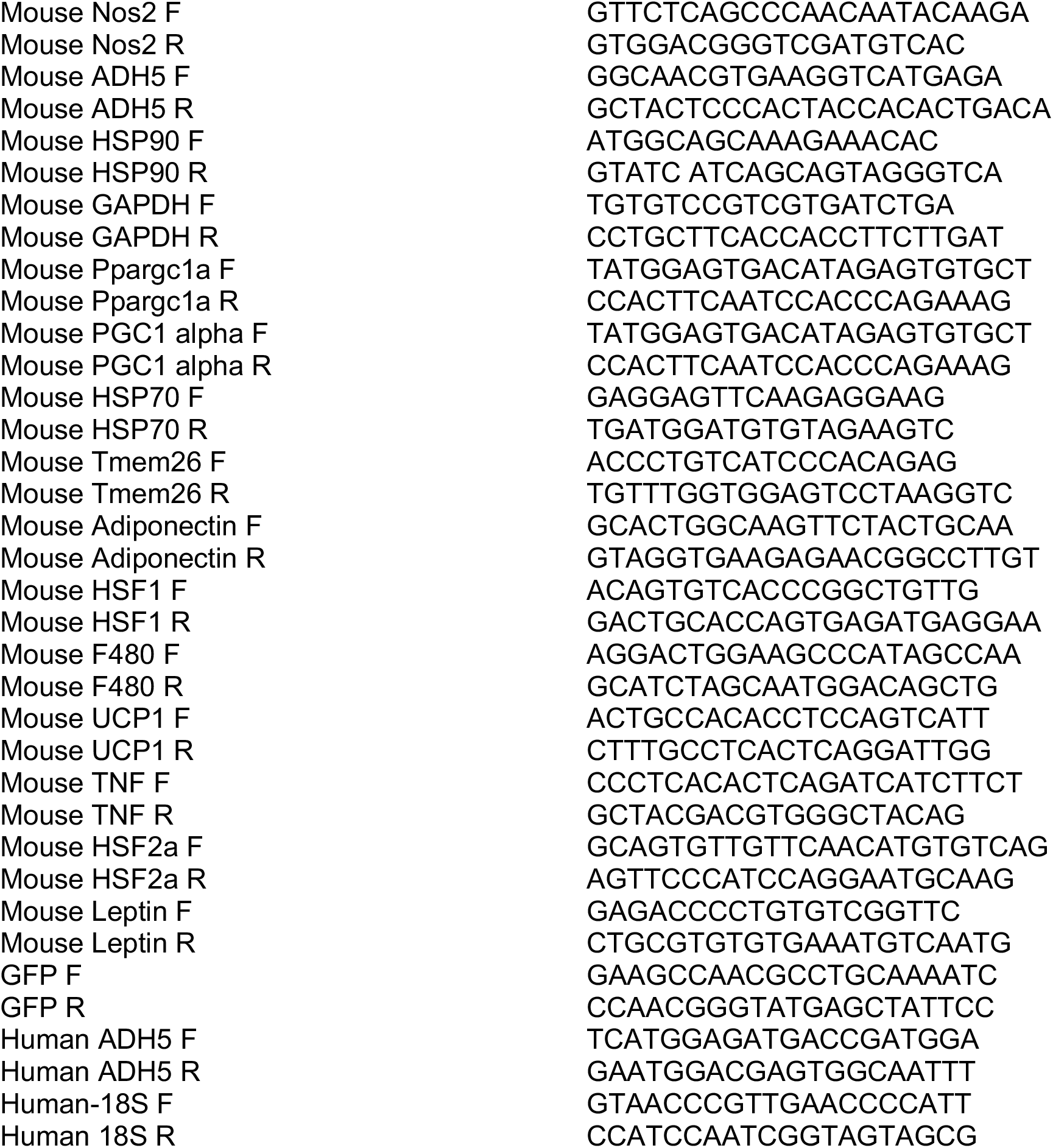

### Western blot Analysis

Proteins were extracted from cells or tissues and subjected to SDS–polyacrylamide gel electrophoresis, as previously described(Yang et al., 2015). Membranes were incubated with anti-ADH5 (Abcam, ab91385), anti-pAKT (Cell Signaling Technology, 4060), anti-AKT (Santa Cruz, SC-8312), anti-UCP1 (Abcam, ab10983), anti-Actin (Santa Cruz, sc-47778) or anti-Tubulin (Cell Signaling Technology, 11H10) at 1:1000 dilution; and then incubated with the appropriate secondary antibody conjugated with horseradish peroxidase (1:5000, Santa Cruz, sc-2005). Signal was detected using the ChemiDoc Touch Imaging System (Bio-Rad), and densitometry analyses of western blot images were performed by using the Image Lab software (Bio-Rad).

### Nuclear fractionation

The nuclear fractions were prepared as described previously (Yang et al., 2015). Briefly, 30μg BAT tissue was homogenized in hypotonic buffer: 250 mM sucrose (Sigma-Aldrich, S7903), 20 mM HEPES, pH 7.5 (Sigma-Aldrich, H3375), 10 mM KCl (Sigma-Aldrich, P9333), 1.5 mM MgCl_2_ (Sigma-Aldrich, 208337), 1 mM EDTA (Sigma-Aldrich, EDS), and 1 mM EGTA (Sigma-Aldrich, E3889) followed by filtering through a 100 μm cell strainer. The cell lysates were centrifuged at 825 g for 20 minutes at 4°C. The pellets were dissolved in NE buffer (20 mM HEPES, pH 7.9, 1.5 mM MgCl_2_, 0.5 M NaCl (Sigma-Aldrich, S7653), 0.2 mM EDTA, 20% glycerol (Sigma-Aldrich, G5516) and passed through a 32G needle. The lysate was further cleared by spinning at 18000 g for 15 minutes at 4°C.

### S-nitrosylation detection

#### In situ detection of S-nitrosylated protein

was performed as described with minor modifications (Thibeault et al., 2010). Briefly, frozen BAT sections were fixed with 3.5 % paraformaldehyde, and then permeabilized with 0.1 % Triton X-100 in PBS containing 1mM EDTA and 0.1mM neocuproine. Biotin-switch assay was performed by first blocking free thiols using HENS buffer containing 20 mM MMTS at room temperature for 30 mins. Then the S-nitrosylated proteins were labeled in HENS buffer with 0.4mM biotin-HPDP and 10mM ascorbic acid for 1hr.Biotinylated proteins were labeled using strepvidin conjugated with Alexa 488. The sections were then subject to immunostaining for UCP1 using anti-UCP1 (1:1000, Abcam, ab10983) and secondary antibodies conjugated to Alexa-568. The images were observed using a Ziess 700 confocal or Leica fluorescence microscope. The images were quantified using ImarisColoc (Bitplane).

#### Detection of S-nitrosylation proteins

Total S-nirosylated proteins in the BAT was detected by S-nitrosylation Western Blot Kit (ThermoFisher Scientific). Briefly, the BAT tissues were lysed in HEN buffer and 500ug protein was used for each sample. Free cysteines were first blocked with MMTS for 20 mins at 50°C. Following precipitation using 4× volume of cold acetone for 60 min at −20 °C, S-nitrosylated cysteines were selectively labeled with iodoTMT reagent in the presence of 20mM sodium ascorbate at room temperature for 2 hrs. Samples were then separated by SDS-PAGE and blotted to membrane for detection. SNO proteins were detected by the anti-TMT antibody (ThermoFisher Scientific).

#### Assessment of SNO of UCP1 by mass spectrometry

100μg recombinant human UCP1 (MyBioSource) was treated with or without of 200μM SNAP for 15 mins in dark at RT. The samples were then blocked with MMTS for 20 mins at 50°C to block free cysteines, and labeled with iodoTMT reagent in the presence of 20mM sodium ascorbate at room temperature for 2 hrs. After acetone precipitation and desalt processes, the samples were subjected to mass spectrometry analysis performed by the Taplin Biological Mass Spectrometry Facility at the Harvard Medical School using an Orbitrap Fusion mass spectrometer. MS/MS data were searched against the Uniprot mouse database using the SEQUEST algorithm.

### ADH5 Activity assay

The GSNOR enzyme activity assay was performed as described previously (Liu et al., 2001). Briefly, 10μg protein lysate was added to 200μl of assay mix (20mM Tris-Hcl, 200μM NADH, and 0.5mM EDTA). The kinetic of GSNO-dependent NADH consumption was measured in the absence or presence of 100μM of GSNO by using a microplate reader (340 nm) at 37°C. The resulting GSNOR activity was expressed as nmol NADH degraded min/ mg protein.

### Oil Red O staining

Murine or human differentiated brown adipocytes were fixed with 4% PFA and stained with 0.3% Oil Red O solution (Sigma-Aldrich, O0625). The images were observed under a Nikon microscope.

### Immunohistochemistry and Immunofluorescence

For immunohistochemistry, iBAT depots were fixed with 4% PFA and sectioned at 5 μm thick, followed by deparaffinization and rehydration processes. Tissue sections were stained using H&E. The images were observed under a Nikon microscope (10x). For F4/80 staining, antigen retrieval was performed using heat, and the slides were then permeabilized with 0.1% Triton X-100 in PBS. Sections were incubated overnight at 4°C with F4/80 antibody (1:100, Cell Signaling, D2S9R) followed by secondary staining for one hour at RT (1:250, Alexa-488, Invitrogen, A11008). Images were taken using a LSM 510 confocal microscope (Carl Zeiss) and analyzed with NIH ImageJ. For ADH5 staining, sections were stained using an anti-ADH5 antibody (1:100, Abcam, ab91385). Histochemical reactions were performed using a Vectastain ABC kit (Vector Laboratories) and Sigma Fast 3,3′-diaminobenzidine (Sigma-Aldrich, D4293) as the substrate. Images were taken by using a Nikon microscope.

### Lipolysis assay

Lipolysis in BAT was measured as described previously (Bartelt et al., 2018). Briefly, BAT depots were excised, cut into three pieces per mouse and incubated in 24-well plates in DMEM with high glucose supplemented with 2% wt/vol FA-free BSA (FA-free BSA DMEM, Sigma). BAT was then incubated for 1 hour at 37 °C before stimulation with 10 μM isoproterenol (ISO, Sigma, 54750-10-6). Samples were taken at the indicated time points, and the supernatant was assayed for free fatty acid (FFA) using a FFA fluorometric kit (Cayman Chemical, 700310). The fatty acid release was normalized to RNA content.

### Metabolic Phenotyping

#### Whole-body energy expenditure and body composition

Whole-body energy expenditure (VO_2_, VCO_2_), food intake, and locomotor activity were monitored using a Comprehensive Lab Animal Monitoring System (CLAMS, Columbus Instruments) at the Fraternal Order of Eagles Metabolic Phenotypic Core. For core temperature measurements, a thermistor was implanted subcutaneously within the abdominal cavity; for BAT temperature recording, a thermistor was implanted between the BAT and the underlying muscle layer in the inter-scapular region prior. The temperature at each tissue was recorded by CLAMS Telemetry Temperature Transmitter system. Body composition was measured by using Bruker Minispecs (LF50).

#### Infrared imaging

Mice were injected with norepinephrine (0.3mg/kg) via i.p. 15 mins after injections, body surface temperature was imaged in fully awake, unrestrained mice using a high-resolution infrared camera (A655sc Thermal Imager; FLIR Systems, Inc.) as described (Zhu et al., 2014).

#### Respirometry of BAT

Freshly isolated iBAT were weighed (∼2mg) and minced in respiration buffer (2.5 mM glucose, 50 μM palmitoryl-L-carnitine hydrochloride, 2.5 mM malate, 120 mM NaCl, 4.5 mM KCl, 0.7 mM Na_2_HPO_4_, 1.5 mM NaH_2_PO_4_ and 0.5 mM MgCl_2_, pH 7.4). High-resolution O_2_ consumption was measured in 2 mL of buffer Z containing 0.5mM EDTA using the OROBOROS Oxygraph-2k (O2k; Oroboros Instruments, Innsbruck, Austria (Tyrrell et al., 2016). All respiration measurements were conducted at 37°C and a working range [O_2_] of ∼350–180 μM. Respiration was measured as follows: 5 mM malate + 10 mM pyruvate followed by 1mM GDP (UCP1) and finally 0.5 mM ADP (complex I OXPHOS substrate). Data shown correspond to subtraction of the oxygen flux observed after guanidine diphosphate (GDP) addition (1μM, for UCP1) or 0.5mM adenosine diphosphate (ADP, for Complex I) from that measured after the addition of 10mM pyruvate + 5mM malate and normalized to baseline flux. The graph shown corresponds to subtraction of the oxygen flux observed after addition of 1μM guanidine diphosphate (GDP) from that measured after addition of 10mM pyruvate + 5mM malate and normalized to baseline flux.

#### Primary brown adipocyte cellular respirometry

Cellular OCR of brown adipocyte cells was determined using a Seahorse XFe96 Analyzer. Adipocytes were plated and differentiated in XF96 cell culture microplates, and differentiated for 6 days. Basal respiration was determined to be the OCR in the presence of substrate alone. The respiratory rate was measured at 37°C in 8 replicates (independent wells). ATP-synthase-independent respiration was determined after addition of 2.5 μM oligomycin, leak respiration was determined after addition of 2.5 μM oligomycin and 100 nM noradrenaline. Maximal respiration was determined after addition of 2 μM FCCP.

#### Glucose tolerance test

Glucose tolerance was tested by intraperitoneal glucose injection (0.8-1.25 g/kg body weight, 50% dextrose, Hospra Inc, 0409-6648-02) (Qian et al., 2018).

### Statistical Analysis

Results are expressed as the mean ± the standard error of the mean (SEM); *n* represents the number of individual mice (biological replicates) or individual experiments (technical replicates) as indicated in the figure legends. We performed the Shapiro-Wilk Normality test in experiments that have a relatively large sample size (n>5) and found that these data pass the normality test (alpha=0.05). Data were further analyzed with two-tailed Student’s and Welch’s *t*-test for two-group comparisons or ANOVA for multiple comparisons. For both One-Way ANOVA and Two-Way ANOVA, Tukey’s post-hoc multiple comparisons were applied as recommended by Prism. In all cases, GraphPad Prism (GraphPad Software Prism 8) was used for the calculations.

